# Clusters of lineage-specific genes are anchored by ZNF274 in repressive perinucleolar compartments

**DOI:** 10.1101/2024.01.04.574183

**Authors:** Martina Begnis, Julien Duc, Sandra Offner, Delphine Grun, Shaoline Sheppard, Olga Rosspopoff, Didier Trono

**Affiliations:** School of Life Sciences, Ecole Polytechnique Fédérale de Lausanne, Lausanne, Switzerland

## Abstract

Long known as the site of ribosome biogenesis, the nucleolus is increasingly recognized for its role in shaping 3D genome organization. Still, the mechanisms governing the targeting of selected regions of the genome to nucleolus-associated domains (NADs) remain enigmatic. Here we reveal the essential role of ZNF274, a SCAN-bearing member of the Krüppel-associated box (KRAB)-containing zinc finger proteins (KZFP) family, in sequestering lineage-specific gene clusters within NADs. Ablation of ZNF274 triggers transcriptional activation across entire genomic neighborhoods – encompassing, among others, protocadherin and KZFP-encoding genes – with loss of repressive chromatin marks, altered 3D genome architecture and *de novo* CTCF binding. Mechanistically, ZNF274 anchors target DNA sequences at the nucleolus and facilitates their compartmentalization via a previously uncharted function of the SCAN domain. Our findings illuminate the mechanisms underlying NADs organization and suggest that perinucleolar entrapment into repressive hubs constrains the activation of tandemly arrayed genes to enable selective expression and modulate cell differentiation programs during development.

## Introduction

The eukaryotic genome is spatially organized to guarantee the orderly execution of transcriptional programs^1,2^. With hundreds of genes switching between active and inactive compartments during development and differentiation^3,4^, nuclear bodies can serve as a scaffold for this reorganization^5,6^. In most nuclei, heterochromatin gathers at the nuclear lamina or around the nucleolus. Whilst lamina-associated domains (LADs) are extensively mapped, the membrane-less nature of the nucleolus has hampered the identification of their nucleolus-associated domains (NADs) counterparts^8,11^. Thus, what is the role of perinucleolar heterochromatin in genome organization and what directs specific gene subsets to this region remain unanswered questions.

The nucleolus is the largest identified nuclear substructure and is best known as the site of ribosome biogenesis. Silent ribosomal DNA (rDNA) repeats condense at the nucleolar periphery together with centromeres and pericentromeric regions. Early investigations into human NADs using biochemical purification of nucleoli revealed an enrichment of specific gene subsets, including protocadherins, immunoglobulins, and olfactory receptors, which are all characterized by their organization in gene clusters, heterochromatinization and cell type-restricted expression^8,12–15^. These findings led to the hypothesis that the nucleolus may act as a specialized compartment for establishing repressive chromatin and regulating dynamic lineage-specific expression programs.

Unraveling the determinants of NADs is critical to shed light on the role of the nucleolus in broad epigenetic regulatory mechanisms and comprehend how nuclear compartments can impact gene expression patterns in development and disease. While a few proteins have been implicated in assisting chromatin interactions with the nucleolus(*16*, *17*), whether they truly serve as direct anchors was not established. Amongst the gene clusters enriched at NADs are *KZFPs*, responsible for transcription factors encoded in hundreds by the human genome. *KZFP* genes are decorated with a unique H3K9me3 chromatin signature deposited over their 3’ end by one of their family member, ZNF274, via the KRAB-mediated recruitment of KAP1/TRIM28 and associated SETDB1^18^. Notably, ZNF274 was shown to have nucleolar targeting ability and computational modeling found it to be a TAD boundary-associated genomic element^19,20^. Despite these intriguing connections linking ZNF274 to the perinucleolar environment, its role in 3D chromatin architecture and gene regulation has so far remained undetermined. The present study reveals that it is responsible for targeting clusters of lineage-specific genes to repressive perinucleolar subdomains.

## Results

### ZNF274 represses lineage-specific genes organized in clusters

To analyze the biological function of ZNF274, we generated homozygous knockout (KO) HEK293T cells by CRISPR/Cas9 genome editing (Figure S1A, S1B). Transcriptome profiling by RNA sequencing (RNA-seq) revealed that almost all (79/80) protein-coding genes with significantly altered mRNA levels in *ZNF274* KO cells were upregulated (Figure 1A and Table S1). 60 of them encoded KZFPs and eight some other C_2_H_2_ zinc finger protein (ZFP). Standing out amongst transcripts upregulated in the absence of ZNF274 were also products of the protocadherin (PCDH) gene family. PCDHs are adhesion molecules predominantly expressed in the nervous system and, like KZFPs, are encoded by genes organized into tandem arrays (α, β, and γ clusters on chromosome 5). We found representatives from all *PCDH* clusters to be significantly more transcribed in *ZNF274* KO cells (Figure 1A). We were struck by the similarities of genomic modularity and regulatory logic displayed by *KZFP* and *PCDH* genes, for which the ground state of most promoters is repressed. Nevertheless, previous profiling of ZNF274 genome-wide binding sites did not reveal an enrichment at promoters of genes deregulated in our KO mutants^21^. To explore the issue further, we reintroduced hemagglutinin (HA)-tagged ZNF274 in KO cells and performed chromatin immunoprecipitation sequencing (ChIP-seq) (Figure S1C). Of over 20’000 significantly enriched sites, most were located in introns or intergenic regions (Figure S1D, S1E and Table S2). We found that ZNF274 especially accumulated over the 3′ end of *KZFP* genes but not on other *ZFP* or *PCDH* genes (Figure 1B, 1C and S1F). Motif analysis of ZNF274-associated loci confirmed that the top scoring motif for ZNF274 (Figure 1D) was present in the ZNF-encoding exons of 331 out of 377 *KZFP* genes (Figure 1E). ZNF274 peaks are ingrained in heterochromatin marked by H3K9me3 and these loci failed at maintaining the silencing mark upon ZNF274 depletion (Figure 1F). Remarkably, loss of H3K9me3 was not restricted to ZNF274 peaks but extended across large portions of the genome including regions located in pericentromeric and subtelomeric domains, where lack of repressive histone modifications was concomitant with an increase in the H3K4me3 activating mark (Figure 1G, 1H, 1I). Prominent alterations in chromatin marks were identified within segments housing *KZFP* gene clusters (Figure 1J, S1G). At the *PCDH* locus, two ZNF274-recruiting regions (R1 and R2) interspersed between *PCDH* clusters underwent substantial shrinkage of H3K9me3-adorned chromatin after ZNF274 loss (Figure 1K). On the contrary, a minority of sequences displayed a gain of H3K9me3 and were enriched in binding sites for other KZFPs, mirroring the functional activation of these repressors (Figure S1H, S1I, S1J). Given the profound crosstalk between chromatin modifications and DNA methylation, we used the enzymatic methyl-seq (EM-seq) method to interrogate the impact of the large-scale H3K9me3 contraction on CpG methylation (mCG). We detected only a minor decline of mCG on the promoter of *KZFP* genes, which are mainly marked by unmethylated DNA in HEK293T cells (Figure S2A). On the other hand, many of the H3K9me3 domains affected by loss of ZNF274 showed an overall increase of mCG level (Figure S2B and S2C). The deposition of mCG was particularly pronounced on *KZFP* and *PCDH* gene bodies, where it correlated with the concurrent accrual of H3K36me3 histone mark, known to recruit the *de novo* methyl-transferase DNMT3b (Figure S2D, S2E, S2F)^22^. Likewise, DNA segments harboring ZNF274 peaks displayed increased mCG levels specifically where H3K9me3 had been lost (Figure S2G).

**Fig. 1.**
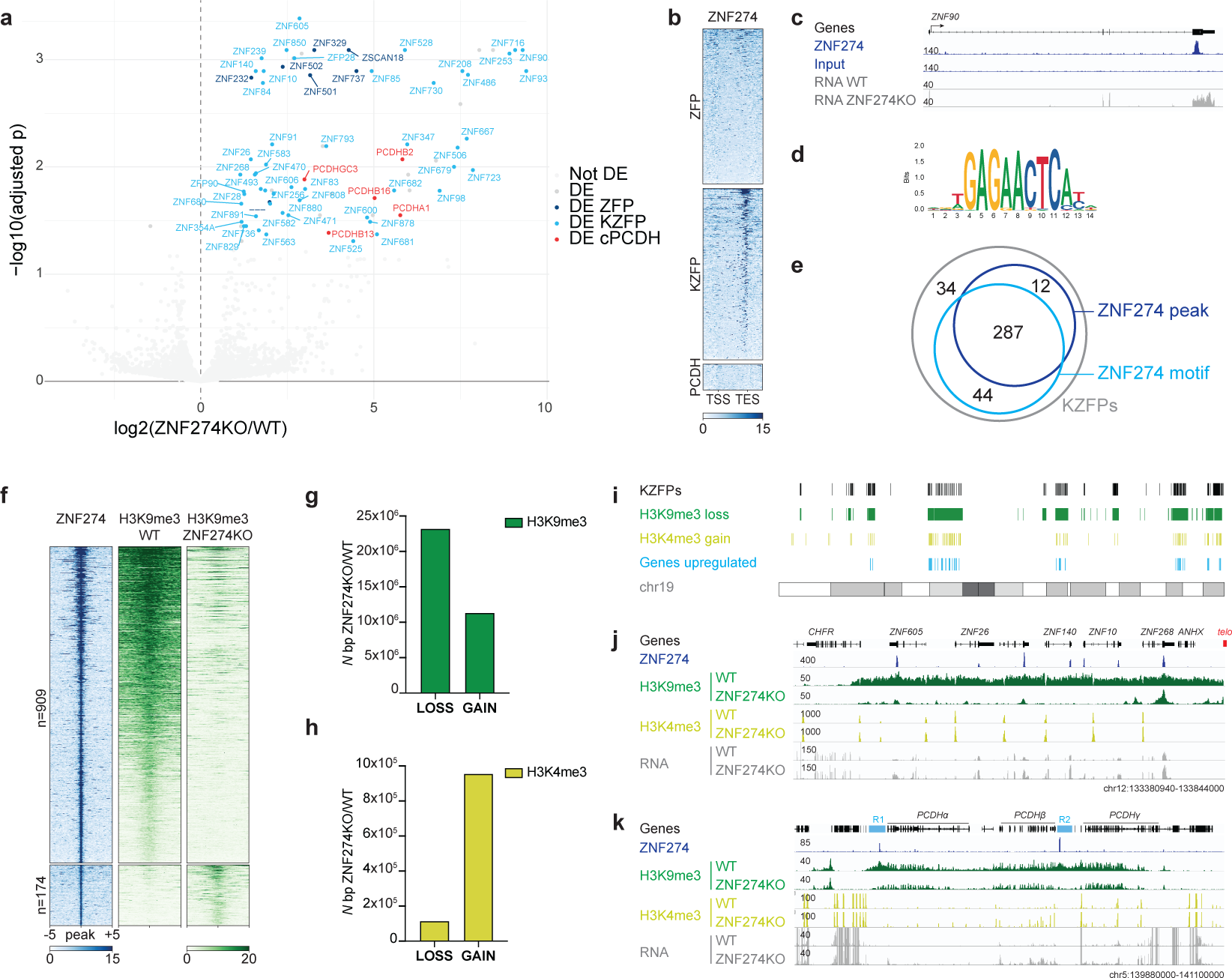
ZNF274 represses lineage-specific genes organized in clusters. **a.** Volcano plot comparing fold change in gene expression for wild-type and *ZNF274* KO HEK293T cells. Representative *ZFP*, *KZFP* and *PCDH* genes are highlighted in blue, light blue and pink. **b.** Heatmap of ZNF274 ChIP-seq enrichment across the body of *ZFP*, *KZFP* and *PCDH* genes (+2 kb from TSS or TES). **c.** IGV browser screenshot of a representative KZFP gene showing tracks for ZNF274 ChIP-seq in *ZNF274* KO cells overexpressing HA-tagged ZNF274, and RNA-seq in wild-type and *ZNF274* KO cells. **d.** Top scoring DNA binding motif identified for ZNF274. **e.** Venn diagram showing overlap of KZFP genes harboring ZNF274 motif and ChIP-seq peaks. **f.** Heatmap of ZNF274 and H3K9me3 ChIP-seq across ZNF274 peaks showing differential enrichment of H3K9me3. Each row represents a 5 kb window centered on peak midpoint, sorted by ZNF274 ChIP signal. **g.** Bar plot depicting significant loss or gain of H3K9me3 in wild-type versus *ZNF274* KO HEK293T cells (FC > 2; adjusted Pvalue < 0.05; n = 4 replicates/group). **h.** Bar plot depicting significant loss or gain of H3K4me3 in wild-type versus *ZNF274* KO HEK293T cells (FC > 2; adjusted Pvalue < 0.05; n = 2 replicates/group). **i.** Karyotype plot of chromosome 19 showing the genomic localization of regions significantly affected upon *ZNF274* knock-out. **j.** IGV browser screenshots showing tracks for ZNF274 ChIP-seq in *ZNF274* KO cells overexpressing HA-tagged ZNF274, H3K9me3 and H3K4me3 ChIP-seq in wild-type and *ZNF274* KO cells across subtelomeric ZFP cluster. The red box indicates the position of telomeric repeats. **k.** IGV browser screenshots showing tracks for ZNF274 ChIP-seq in *ZNF274* KO cells overexpressing HA-tagged ZNF274, H3K9me3 and H3K4me3 ChIP-seq in wild-type and *ZNF274* KO cells across PCDH. ‘R1’ and ‘R2’ indicate the tracts of H3K9me3 chromatin undergoing substantial shrinkage after ZNF274 loss.

The data presented so far suggest that ZNF274-mediated heterochromatin formation is the main mechanism securing robust transcriptional silencing of *KZFP* and *PCDH* gene clusters. Because of the neuron-specific expression of *PCDHs*, we decided to confirm the functional role of ZNF274 in regulating these genes in human neural progenitor cells (NPC) derived from embryonic stem cells (ESC) (Figure S3A, S3B). Genes encoding 28 out of 56 isoforms of PCDH were upregulated in NPC cultures generated from homozygous *ZNF274* KO ESC (Figure 2A and Table S3). We also observed the upregulation of most *KZFP* genes in NPCs (Figure S3C), as well as other tandemly arrayed genes that have lineage-specific functions, such as the kallikrein (*KLK*), pregnancy-specific glycoprotein (*PSG*), keratin (*KRT*) and NOD-like receptor protein (*NLRP*) families (Table S3). Genes downregulated in KO cells were particularly enriched for Gene Ontology terms related to neuron differentiation and brain development (Figure S3D), suggesting that ZNF274 plays a critical role in these processes. Using Cut&Tag, we verified that H3K9me3 levels were globally reduced in KO NPCs, similar to what observed in HEK293T cells (Figure 2B and S3E). Notably, the R1 and R2 loci in the *PCDH* locus lost H3K9me3 in KO cells (Figure 2C), as did all the gene families just listed as upregulated in this setting (Figure S3F-H). In particular, *ZNF274* KO caused loss of H3K9me3 on the Prader–Willi syndrome (PWS)-associated *SNORD116* cluster, known to be epigenetically regulated by ZNF274, stressing the particular relevance of ZNF274 function in the neuronal context (Figure S3I)^23,24^. Conversely, the large *HIST1* cluster on chromosome 6 containing 55 histone genes and localizing within transcriptionally active nuclear bodies (histone locus bodies) was unperturbed (Figure S3J)^25^. We conclude that binding of ZNF274 serves as the pivotal catalyst for the assembly of heterochromatin that permeates genes organized in clusters, thereby thwarting any spurious activation of lineage-restricted expression programs.

**Fig. 2.**
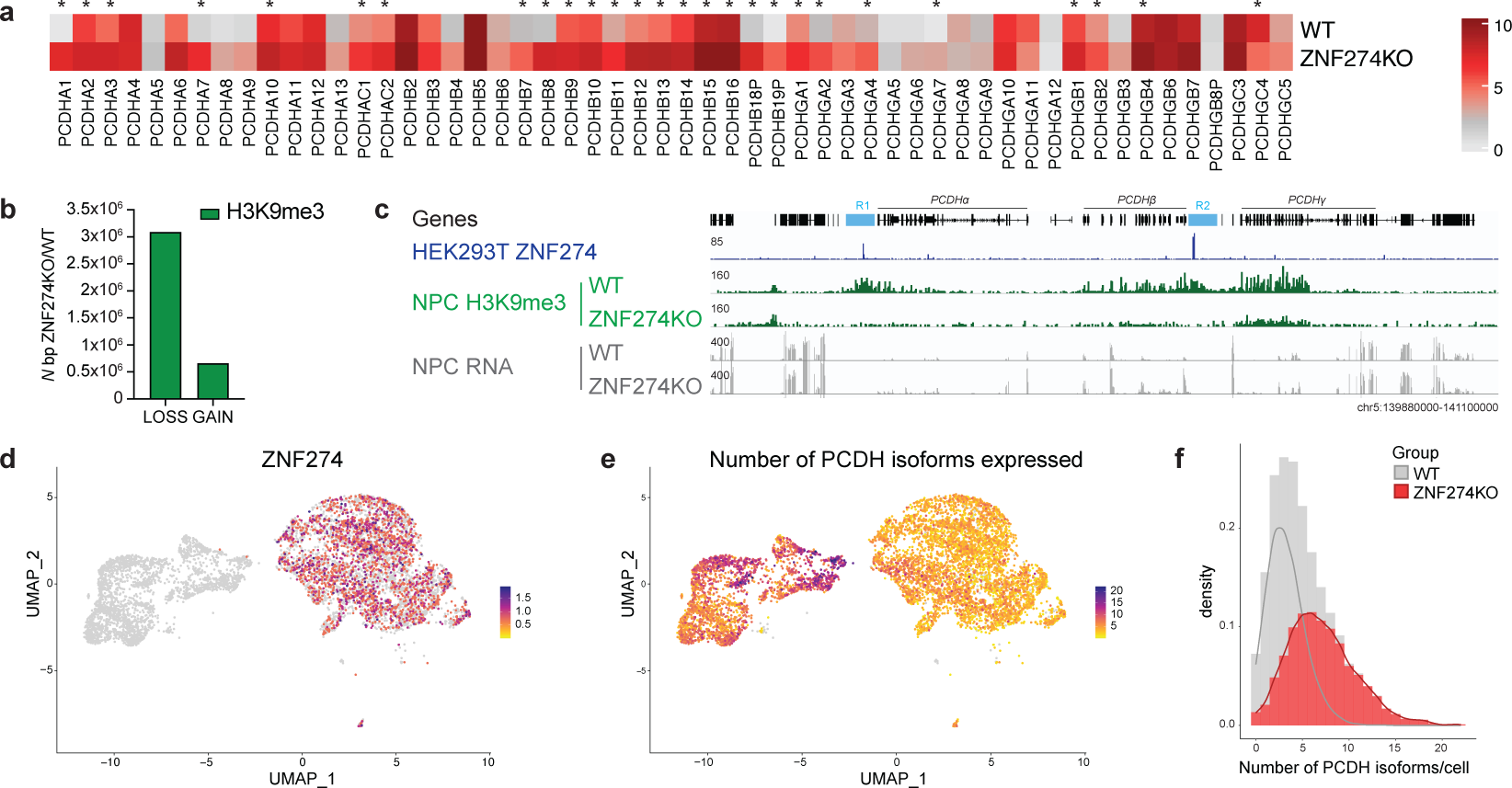
ZNF274 is required for selective and combinatorial expression of *PCDH* genes. **a.** Expressed *PCDH* genes in wild-type versus *ZNF274* KO NPCs. Data are shown as the average of *n* = 2 parental cell lines. Asterisks indicate genes whose expression significantly changes in wild-type versus *ZNF274* KO NPC cells. **b.** Bar plot depicting significant loss or gain of H3K9me3 in wild-type versus *ZNF274* KO NPCs (FC > 2; adjusted Pvalue < 0.05; n = 4 replicates/group). **c.** IGV browser screenshot showing tracks for H3K9me3 Cut&Tag and RNA-seq ChIP-seq in wild-type and *ZNF274* KO NPCs across *PCDH* gene cluster. **d.** UMAP visualization of wild-type and *ZNF274* KO NPCs obtained through 10X 5’-end single-cell sequencing. The UMAP is colored by expression of *ZNF274* gene. **e.** The same UMAP as in (A), colored by expression of number of *PCDH* genes. **f.** Density plot representing the distribution of cells expressing varying numbers of PCDH isoforms either in wild-type and *ZNF274* KO condition. The superimposed curves represent kernel density estimate for each condition.

### ZNF274 is required for selective and combinatorial expression of tandemly arrayed genes

The combinatorial expression of different PCDH isoforms is crucial for proper dendritic arborization because it provides neurons with a unique barcode to distinguish self from non-self^26^. While the mechanisms involved in *PCDH* stochastic promoter choice are difficult to be studied *in vivo* because each neuron expresses at low levels a distinct repertoire of PCDH alternate exons, we could clearly detect a strong gain of expression of PCDH isoforms in our experiments. To test if individual cells expressed an increased number of PCDH isoforms rather than higher levels of individual ones, we performed single-cell RNA-seq of wild-type and KO NPCs (Figure 2D). This revealed that a larger proportion of *PCDH* isoforms were transcribed by each cell upon loss of *ZNF274*, in violation of the ‘one isoform per neuron’ rule that safeguards neuronal self-avoidance (Figure 2E and 2F). In parallel, we found many more *KZFP* genes to be actively transcribed in individual *ZNF274* KO cells (Figure S4A and S4B). Control regions such as the *HIST1* cluster on chromosome 16 maintained similar expression levels in wild-type and KO conditions (Figure S5C and S5D), while in contrast we confirmed the activation of other clustered gene families regulated by ZNF274 (Figure S4E and S4F). Collectively, these findings strongly suggest that ZNF274-dependent repressive mechanisms play an essential role in constraining the selective activation of tandemly arrayed genes, so that loss of ZNF274 activity precludes the fine-tuning of isoform diversity that allows cells to achieve phenotypic and functional diversification during embryogenesis and differentiation.

### ZNF274 occupancy hampers binding of CTCF and promotes genome compartmentalization

The replacement of repressive by activating histone marks upon *ZNF274* KO suggested a widespread recruitment of transcription factors altering enhancer-promoter communication. At the *PCDH* locus, it is well documented that transcriptional activation requires the formation of CTCF-driven DNA looping between gene promoters and downstream enhancers^27–30^. To this end, we performed ChIP-seq analysis of CTCF in HEK293T cells. Newly gained CTCF peaks were found on *PCDH* promoters in the absence of ZNF274 (Figure 3A); likewise, CTCF signal increased across *KZFP* clusters (Figure 3B). In general, regions that experienced a reduction in H3K9me3 showed a mirroring rise in CTCF, whereas in areas that gained H3K9me3, CTCF occupancy decreased (Figure 3C and 3D). 99,4% (173/174) of sequences with significantly altered CTCF recruitment represented enhanced (including *de novo*) peaks (Figure 3E) and were enriched in the vicinity of ZNF274-controlled genes (Figure 3F), with 136/174 containing a CTCF motif (JASPAR MA0139.1, p-value < 0.00005). Because the two proteins often co-localize at CTCF barriers^31^, we verified that the cohesin subunit RAD21 was also recruited to newly gained CTCF sites in KO cells while remaining unchanged at control regions (Figure 3G). The mutual exclusivity between ZNF274 and CTCF occupancy led us to investigate whether the absence of ZNF274 would perturb chromatin spatial organization. We performed Hi-C for three wild-type and three KO HEK293T cell clones and pooled the data after verifying the high quality of the triplicates. Overall chromosomal interaction landscapes were very similar between wild-type and KO cells, but chromosome 19, which is where in human most *KZFP* clusters are found^32^, displayed a striking decrease in the number of far-*cis* contacts (>10 Mb) (Figure S5A). Genome partitioning of chromosome 19 into A and B compartments was exceptionally affected in KO cells, indicating that the segregation between *KZFP*-containing compartments was less strict in this setting (Figure 3H and 3I). Other *KZFP* clusters located on different chromosomes showed similar defects; elsewhere, the interaction landscape was unaltered (Figure S5B-E). Analysis of the Hi-C data revealed a weakening of insulation of the *PCDH* locus, concomitant with the appearance or strengthening of architectural “stripes” along the α, β, and γ clusters (Figure 3J), a feature that has been associated with the activity of cohesins in the assembly of enhancer-promoter complexes^33^. Local rewiring of chromatin interactions at the *PCDH* locus was further supported by a series of UMI-4C experiments using contact profiling baits targeting ZNF274 peaks in R1 and R2 as well as the annotated enhancers HS5-1 and HS16 (Figure S6A). UMI-4C data confirmed the changes observed in our Hi-C maps and yielded additional observations. Not only the relative contact strength between enhancers and *PCDH* alternate exons was increased in KO compared to wild-type cells, but we could also detect a reduction in long-range contacts between R1 and R2 in the absence of ZNF274. R1 and R2 correspond to robust SETDB1-sensitive H3K9me3 peaks that in mice have been shown to form repressive loop bundles regulating the *PCDH* locus^27^. In human, R1 polymorphism has been associated with an increased risk of schizophrenia (chr5: 140,023,664–140,222,664) by the Psychiatric Genomics Consortium^34^. R1 carries the noncoding SNP rs11189671 about 70bp from the ZNF274 identified motif, whose dosage has been proven to correlate the expression of multiple *PCDH* genes in adult frontal cortex^35^. Using a GFP reporter system, we confirmed that R1 acts as a ZNF274-dependent *cis*-repressor and that the risk SNP rs111896713 partially dampens its influence (Figure S6B and S6C). Therefore, ZNF274-controlled R1 and R2 serve as key organizers for local chromosomal conformations at the *PCDH* locus and exert repressive effects on gene expression. Unexpectedly, we discovered long-range DNA contacts between *PCDH* and two *KZFP* clusters located ∼10Mb and ∼40Mb away respectively (Figure 3L and S5F). These contacts, which depend on ZNF274, imply the presence of a shared nuclear subcompartment where these genes potentially segregate for co-regulation. The formation of ZNF274-dependent multi-cluster hubs is a conserved feature across different cell types, with the same type of *trans*-interactions between *KZFP* and *PCDH* genes detected in both HEK293T cells and NPCs (Figure S7A-D). Taken together, our data demonstrate that binding of ZNF274 instigates rearrangements in 3D genome architecture that are tailored to segregate lineage-specific genes into repressive compartments impermeable to CTCF occupancy.

**Fig. 3.**
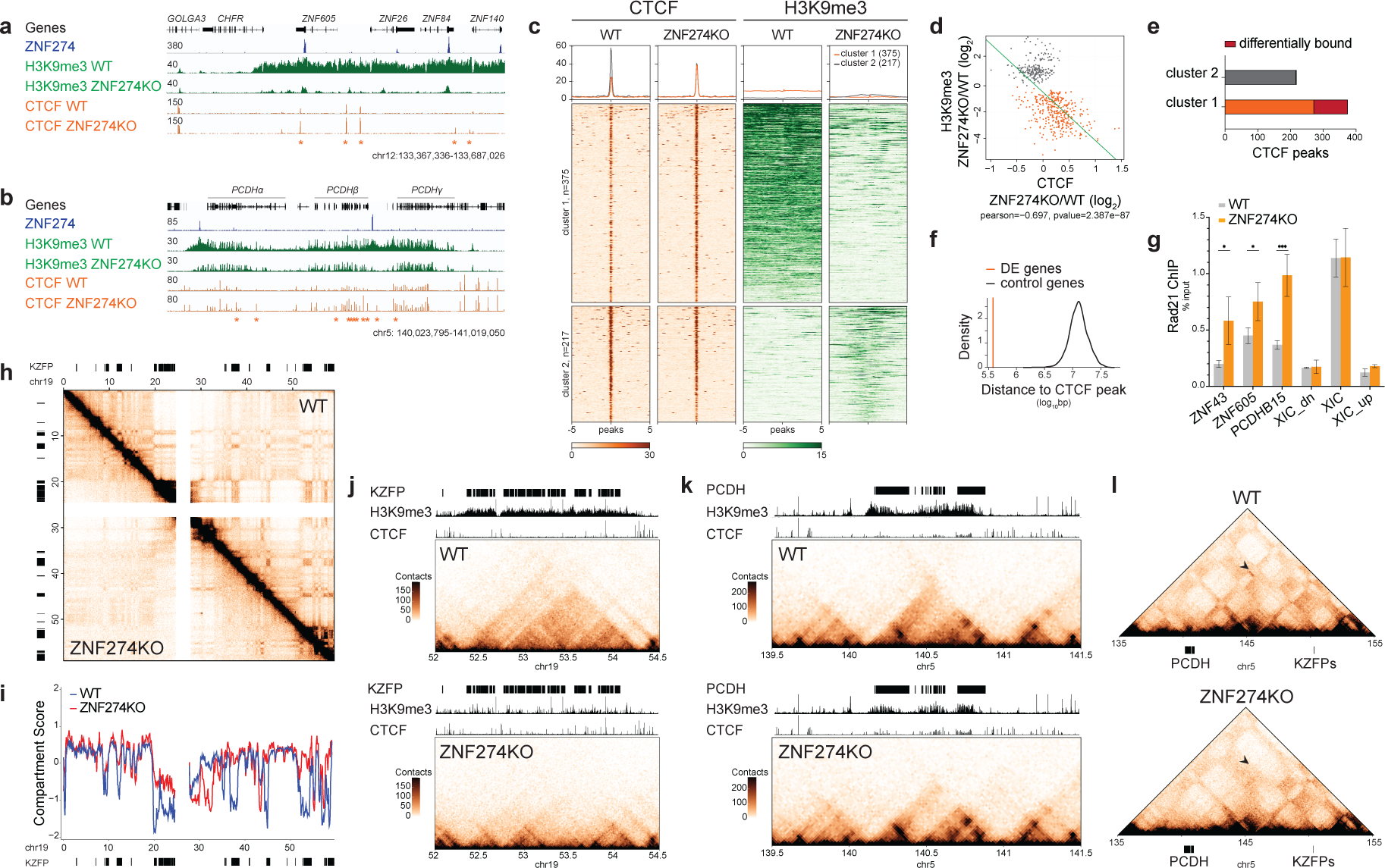
ZNF274 occupancy hampers binding of CTCF and promotes genome compartmentalization. **a.** IGV browser screenshot showing tracks for ZNF274 ChIP-seq in *ZNF274* KO cells overexpressing HA-tagged ZNF274, H3K9me3 and CTCF ChIP-seq in wild-type and *ZNF274* KO cells across *ZFP* clusters (* adjusted P-value<0.1, FC>1.5). **b.** IGV browser screenshot showing tracks for ZNF274 ChIP-seq in *ZNF274* KO cells overexpressing HA-tagged ZNF274, H3K9me3 and CTCF ChIP-seq in wild-type and *ZNF274* KO cells across PCDH (* adjusted P-value<0.1, FC>1.5). **c.** Heatmap and quantification of CTCF and H3K9me3 ChIP-seq enrichment across all CTCF peaks lying within hypo- or hyper-methylated H3K9me3 regions. Each row represents a 5 kb window centered on CTCF peak midpoint. **d.** Scatterplot displaying the inverse correlation between H3K9me3 and CTCF enrichment in wild-type versus *ZNF274* KO cells. Pearson correlation coefficient was calculated to measure the relationship between the two variables. **e.** Bar plots depicting significant loss or gain of CTCF in wild-type versus *ZNF274* KO HEK293T cells (FC > 1.5; adjusted Pvalue < 0.1; n = 3 replicates/group). **f.** Density plot showing average distance of significantly different CTCF peaks from differentially expressed genes in *ZNF274* KO cells compared to a ‘random’ set of control genes. **g.** ChIP-qPCR showing enrichment of RAD21 on significantly enriched CTCF peaks in *ZNF274* KO HEK293T cells. XIC = X-inactivation center; XIC_dn and XIC_up are negative controls respectively downstream an upstream of XIC. **h.** Normalized Hi-C matrix at 100kb resolution of chromosome 19 displaying contact frequencies in wild-type (upper triangle) and *ZNF274* KO (lower triangle) HEK293T cells. **i.** Compartment scores of chromosome 19 show changes in segregation into A (>0) and B (<0) compartments. **j.** Normalized Hi-C maps at 20kb resolution for a KZFP gene cluster in wild-type and *ZNF274* HEK293T cells. On the top are reported tracks visualizing KZFP genes (black bars) and ChIP-seq signal of H3K9me3 and CTCF for each relative condition. **k.** Normalized Hi-C maps at 20kb resolution for the PCDH gene cluster in wild-type and *ZNF274* KO HEK293T cells. On the top are reported tracks visualizing PCDH genes (black bars) and ChIP-seq signal of H3K9me3 and CTCF for each relative condition. **l.** Pyramid plot at 100kb resolution showing long-range contacts (black arrow) between PCDH and KZFP on chromosome 5 in wild-type and *ZNF274* KO HEK293T cells.

### The SCAN domain is responsible for bringing ZNF274 to the nucleolus and promoting the formation of repressive compartments

Next we asked how ZNF274 ablation could trigger such alterations in chromosomal conformations. Previous affinity purification and mass spectrometry (AP-MS) experiments identified some nucleolar proteins in the ZNF274 interactome, which supported earlier studies linking ZNF274 to this nuclear subcompartment^19,36^. We performed proximity biotinylation experiments targeting the TurboID enzyme fused to Protein A (ProtA-TurboID) to HA-tagged ZNF274 with an HA-specific antibody, followed by streptavidin-based affinity enrichment and quantitative mass spectrometry (Figure 4A and Table S4)^37^. Besides KAP1 and two other TRIM proteins (TRIM24 and TRIM33) also part of the Transcriptional Intermediary Factor 1 (TIF1) family, hits included several chromatin-associated proteins such as the CHD4/NuRD complex, validating our experimental approach. Most importantly, 16 out of 127 significant ZNF274 interactors were proteins with known nucleolar localization, an enrichment that has never been detected in the interactome of other KZFPs. ZNF274 contains the auxiliary SCAN domain in between two repressive KRAB motifs (Figure 4B). To characterize the specificity of ZNF274 interactions, we overexpressed mutated versions of ZNF274 bearing deletions either of the two KRAB domains (ΔKRAB-ZNF274) or the SCAN domain (ΔSCAN-ZNF274) in KO cells (Figure 4B). Confocal imaging of full-length ZNF274 revealed an exclusive localization in the nucleus with a granular pattern reinforced at the nucleolar rim (Figure 4C); ΔKRAB-ZNF274 molecules were more prominently concentrated at nucleoli, while the SCAN-devoid derivative was excluded from these nuclear sub-compartments (Figure 4C). ProtA-TurboID experiments using ΔSCAN- or ΔKRAB-ZNF274 mutants as baits confirmed two markedly distinct interactomes, with nucleolus-associated proteins enriched in ΔKRAB-ZNF274 extracts and depleted in their ΔSCAN-ZNF274 counterparts (Figure 4D and Table S5). Hence, protein interactions mediated by the SCAN domain are important for ZNF274 nucleolar anchoring. Overexpression of full-length ZNF274 in KO cells was sufficient to restore H3K9me3 formation over target DNA, whereas ΔKRAB-ZNF274 was ineffective (Figure 4E and S7A-B). Complementation of KO cells with ΔSCAN-ZNF274 led to a partial rescue, with some nucleation of H3K9me3 occurring on ZNF274 peaks but a defect in spreading of this histone mark. We confirmed by HA ChIP-seq that both ΔKRAB-ZNF274 and ΔSCAN-ZNF274 constructs retained the ability to bind DNA, although the average height of the peaks was somehow reduced with respect to full-length ZNF274 (Figure S7C). Of note, neither mutant induced any ectopic DNA binding (data not shown). We recorded similar defects in repression using the GFP reporter assay, where the silencing capacity of ZNF274 was in part reduced by the absence of the SCAN domain (Figure S7D and S7E). Compared to punctual nucleation, spreading of heterochromatin requires the ability of transcription factors to stabilize their contacts via homo- or heterodimerization, a known attribute of the SCAN domain^38^. Amongst over 70 SCAN-containing proteins encoded in the human genome^39,40^, only ZNF24 was enriched in the ZNF274 interactome (Figure 4A). Still, neither ChIP-seq signals nor cellular localization of ZNF24 suggest a co-localization with ZNF274^40,41^. We postulated that ZNF274 would function as a homodimer accumulating around nucleoli and bringing bound loci in spatial proximity. We performed co-IP experiments by co-expressing HA- and Flag-tagged derivatives of ZNF274, which confirmed that the protein can homodimerize (Figure 4F). Removal of the SCAN domain abolished this property, whereas the KRAB-deleted mutant failed to recruit KAP1 but maintained its dimerization capacity. Last, using the Hi-C technique, we investigated if expression of ZNF274 or ΔSCAN-ZNF274 constructs could reestablish long-range chromosomal contacts in KO cells. Although ectopic reintroduction of ZNF274 was not sufficient to reinstall genome-wide long-range plaid patterns as in wild-type cells (Figure S9A and S9B), we could detect the reinstatement of some interactions between the *PCDH* locus and some *KZFP* genes ∼10Mb distant (Figure 4G and S9C). On the contrary, expression of ΔSCAN-ZNF274 mutant could not foster such contact formation, even if the H3K9me3 profiles looked analogous in the two conditions (Figure S9D and S9E). In conclusion, our results demonstrate that ZNF274 can be tethered to the perinucleolar environment via SCAN-mediated interactions that instruct the subcellular localization of the protein and contribute to the formation of long-range contacts between repressed gene clusters.

**Fig. 4.**
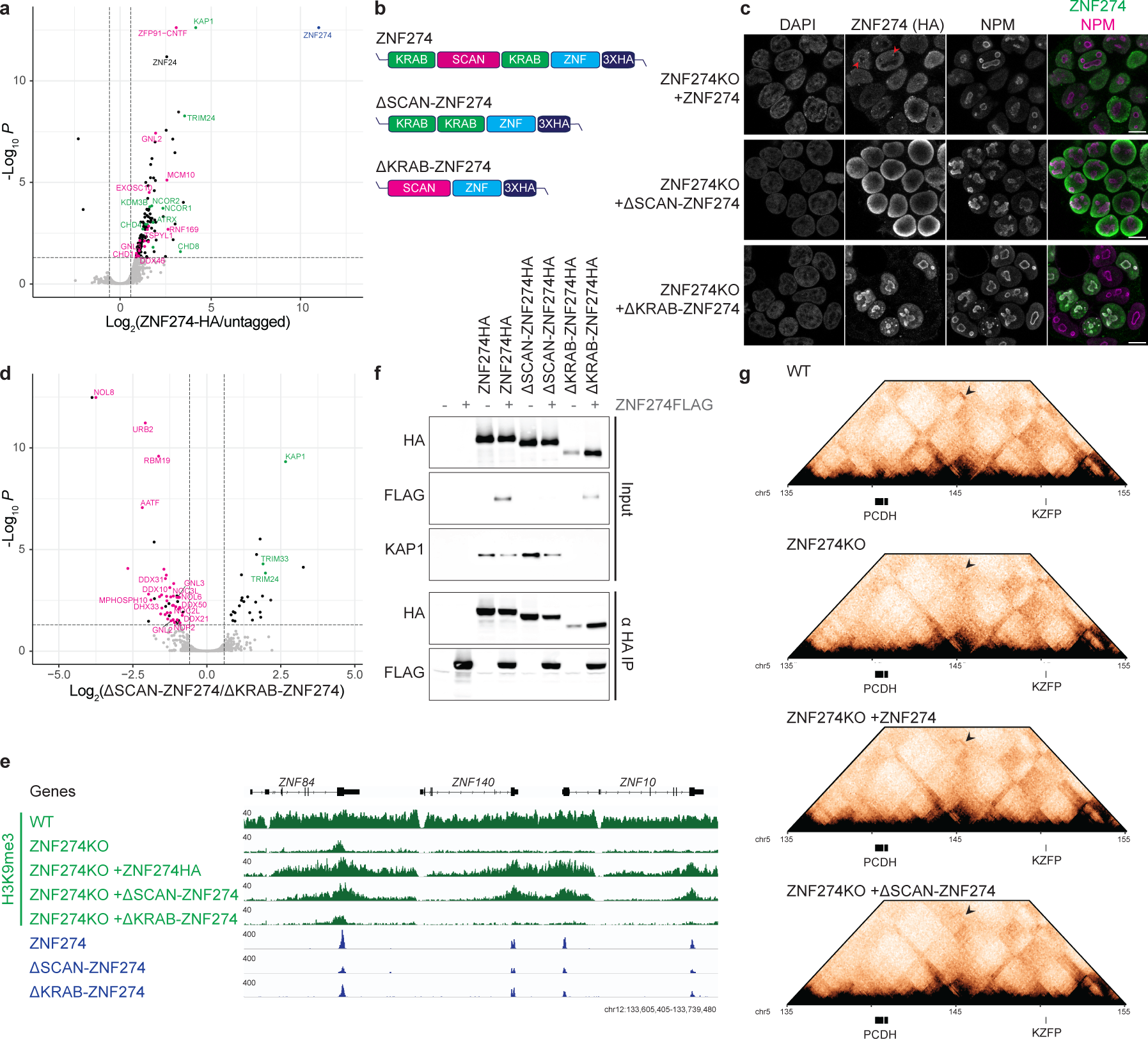
The SCAN domain is responsible for bringing ZNF274 to the nucleolus and promoting the formation of repressive compartments. **a.** Volcano plot of mass spectrometry analysis of biotinylated proteins after targeting ProteinA-TurboID to ZNF274-HA in HEK293T cells (n=3 replicates/condition). A selection of significant interactors are highlighted. Green dots indicate silencer complexes, pink dots indicates proteins known to be localized at nucleoli, blue dot indicates ZNF274 (bait). **b.** Schematic representation of ZNF274 domains and relative mutants. **c.** Representative confocal images of doxycycline induced ZNF274-HA, ΔKRAB-ZNF274-HA, ΔSCAN-ZNF274-HA, the nucleolar marker B23 (NPM) protein and DAPI in *ZNF274* KO HEK293T cells. **d.** Volcano plot of mass spectrometry analysis showing different enrichment of biotinylated proteins after targeting ΔSCAN-ZNF274-HA or ΔKRAB-ZNF274-HA in HEK293T cells (n=4 replicates/condition). Color-coding of interactors is the same as in 4A. **e.** Representative IGV browser screenshot for a KZFP gene cluster showing tracks for H3K9me3 ChIP-seq in wild-type and *ZNF274* KO HEK293T cells overexpressing doxycycline induced ZNF274-HA, ΔSCAN-ZNF274-HA, ΔKRAB-ZNF274-HA, as well as HA ChIP-seq for the same constructs expressed in *ZNF274* KO HEK293T cells. **f.** Co-immunoprecipitation (Co-IP) experiments detecting by western blot HA-tagged ZNF274 proteins, Flag-tagged ZNF274 proteins or KAP1 in cell lysates from HEK293T cells co-transfected with ZNF274 constructs. **g.** Normalized Hi-C pyramid plot at 100kb resolution showing rescue of long-range contacts (black arrow) between PCDH and KZFP gene clusters on chromosome 5 upon overexpression of ZNF274-HA but not ΔSCAN-ZNF274-HA in KO cells.

### ZNF274 tethers bound DNA to nucleoli

A fundamental question raised by the above results is whether ZNF274 directly organizes repressive hubs close to nucleoli. Many of the gene families regulated by ZNF274 are recognized building blocks of NADs^11,12^. We found that about 44% of regions H3K9 hypomethylated in KO HEK293T cells correspond to NAD-associated sequences formerly identified in HeLa cells (Figure 5A, 5B). To probe our hypothesis, we recovered reads from our Hi-C maps containing contacts with ribosomal DNA (rDNA), which is a major nucleolar component. Comparative analysis of inter-chromosomal interactions revealed that contacts between rDNA and ZNF274-targeted DNA were more frequent in wild-type than KO cells (Figure 5C and S10A, S10B). Quantification of rDNA contacts with *KZFP* or *PCDH* genes confirmed a reduced frequency of interactions in the KO condition (Figure 5D). To ascertain directly whether ZNF274 links chromatin to the perinucleolar environment, we performed DNA fluorescence in situ hybridization (FISH) combined with immunofluorescence for nucleolin. Using a probe spanning one ZNF274-regulated *KZFP* cluster, we confirmed that at least one allele (out of the three total) frequently contacts the nucleolus in wild-type cells (Figure 5E). This association was reduced in KO cells, whereas nucleolar localization of the 5S rDNA control region on chromosome 1 stayed unaltered (Figure S10C). These results confirm that ZNF274 instates a comprehensive genome compartmentalization by positioning bound DNA near the perinucleolar environment to regulate accessibility and prevent unintended activation of gene clusters with lineage-specific functions (Figure 5F).

**Fig. 5.**
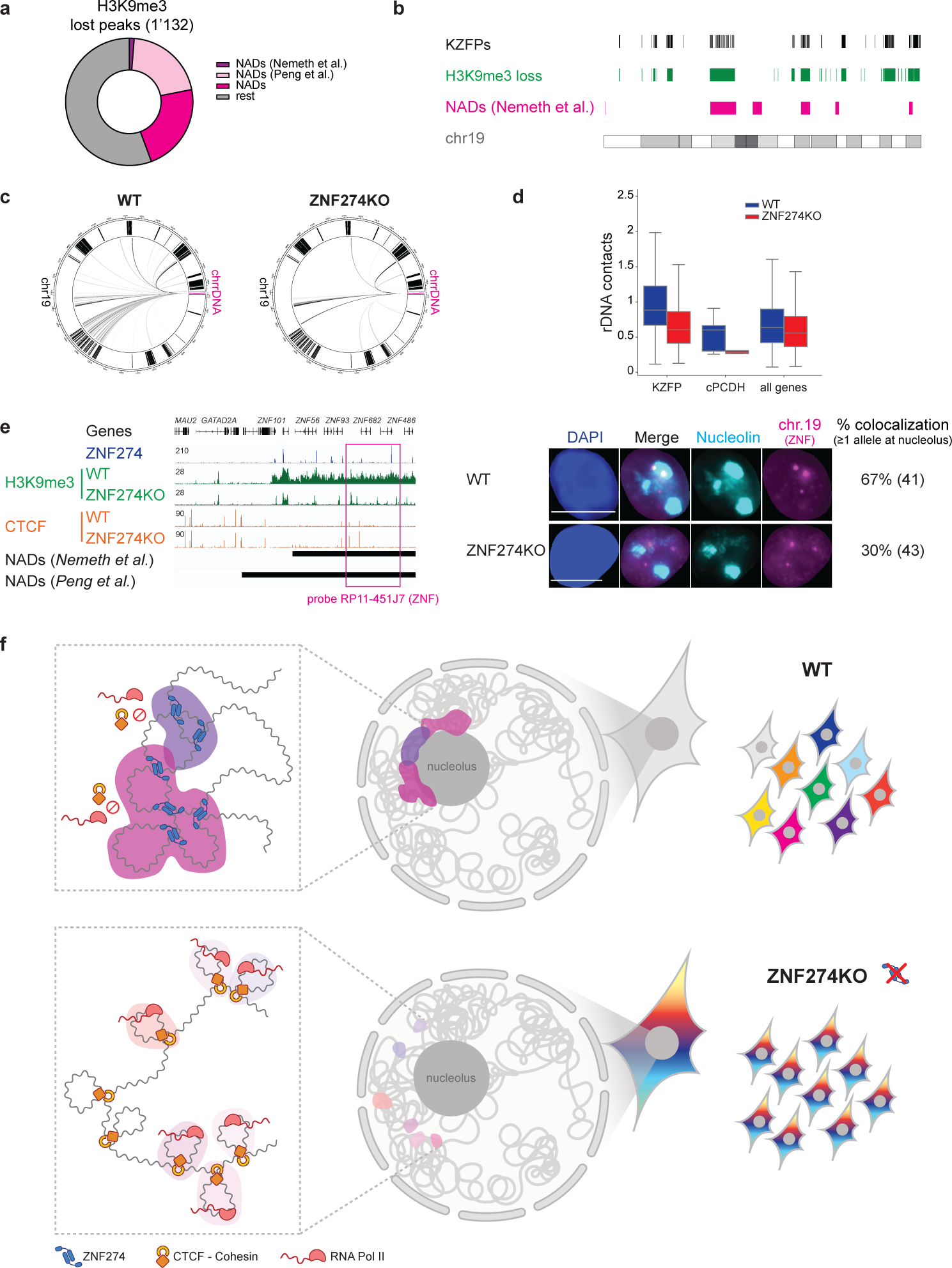
ZNF274 tethers bound DNA to nucleoli. **a.** Doughnut chart displaying the H3K9me3 depleted regions in *ZNF274* KO cells annotated as NADs in previous publications (Nemeth et al., 2010; Peng et al., 2023). **b.** Karyotype plot of chromosome 19 showing the co-localization of H3K9 hypomethylated regions in *ZNF274* KO HEK293T cells and previously annotated NADs in HeLa cells. **c.** Circos plot representation of rDNA contacts with reads mapped on chromosome 19 in wild-type and *ZNF274* KO HEK293T cells using Hi-C matrixes at 100kb resolution. Black straight lines in the outer circle represent *KZFP* genes. **d.** Box-plots (center line, median; box limits, the 25th and 75th percentiles) of trans interactions between rDNA sequences and indicated set of genes (*KZFP*, *PCDH* and all protein-coding genes (∼20’000)) in wild-type and *ZNF274* KO cells. **e.** Left panel: IGV browser screenshot showing localization of the probe used for DNA-FISH. Right panel: 2D ‘maximum projection’ example images from immunofluorescences for nucleolin (light blue) combined with the corresponding DNA-FISH for ZNF cluster on chr19 (pink) and DAPI, with quantification of the number of cells displaying at least one DNA-FISH probe signal contacting the nucleolus in wild-type and *ZNF274* KO HEK293T cells. **f.** Model. (Top) Nucleolar tethering of ZNF274-bound clusters is functional to establish silencing of extended genomic regions and segregate them in repressive domains (in dark pink) that modulate access of CTCF and cohesin complexes to enable selective gene expression. (Bottom) Ablation of ZNF274 triggers loss of repressive chromatin marks, *de novo* CTCF binding and altered 3D spatial organization of the same genomic neighborhoods (in light pink), thus precluding the cell-specific fine-tuning of isoform diversity. The figure panel was created with BioRender.com.

## Discussion

The nucleolus is a central component of nuclear architecture; yet the molecular mechanisms behind this organization have remained elusive thus far. Here we uncovered a previously undescribed mechanism of epigenome regulation for tandemly arrayed genes. Our data suggest a model whereby ZNF274, by targeting specific chromosomal regions near the perinucleolar environment, acts as a guardian against the uncontrolled activation of gene family clusters instead of selective promoter choices (Figure 5F). This is reminiscent of super-repression model proposed by Emile Zuckerkand in 1997, where genomic elements sharing analogous molecular regulatory features meet in the 3D space to spread chromatin modifications in *trans*^42^. It follows that the establishment of sectorial repression must occur prior to lineage-specific transcriptional choices and its long-term stability be ensured by the constitutive production of ZNF274. Supporting this model, ZNF274 is expressed starting at very early embryogenesis and then detected in most tissues. The *ZNF274* gene is also intolerant to loss-of-function mutations, with only a small number of heterozygous individuals (∼20) having been found in large-scale genetic surveys (https://gnomad.broadinstitute.org/). In mice, homozygous mutants of the *ZNF274* ortholog *Zfp110/Nrif1* on a C57BL/6NJCrl background die at embryonic day 12^43^. On the basis of the data presented here, it is reasonable to speculate that loss of ZNF274 is very damaging for human development or even likely embryonic lethal. Furthermore, connections with PWS and schizophrenia pathogenesis suggest that severe intellectual impairments may arise from the dysregulation of ZNF274 expression. Thus, the implications of our findings extend beyond the mechanisms regulating spatial genome organization and hint to a biochemical link between the structural role of NADs and healthy human development.

## ACKNOWLEDGMENTS

We thank Annamaria Kauzlaric, Laia Simo-Riudalbas, Giorgia Brambilla Pisoni, Chase Bolt, and all the members of the Trono laboratory for their technical support and scientific advice; the EPFL Proteomics Core Facility (PCF), the Bioimaging and Optics Platform (BIOP), the Flow Cytometry Core Facility (FCCF) and Gene Expression Core Facility (GECF) for the use of their services and technical assistance. This study was supported by grants from the European Research Council (KRABnKAP, no. 268721; Transpos-X, no. 694658), the Swiss National Science Foundation (310030_192613 and 310030_188803) and the Aclon Foundation to DT.

## AUTHOR CONTRIBUTIONS

M.B. and D.T. conceived the study and designed experiments; M.B. performed the wet experiments with the support of S.O. and O.R; J.D., D.G., S.S., and M.B. conducted the bioinformatics analyses; M.B. and D.T. wrote the manuscript, with review and corrections by all authors.

## COMPETING INTERESTS

M.B. and D.T. are inventors on an international patent application submitted by the École Polytechnique Fédérale de Lausanne for the exploitation of the SCAN domain in genome-editing technologies (European Patent Application EP 23183405.2).

The remaining authors declare no competing interests.

## DATA AVAILABILITY

All data are available in the main text or the supplementary materials, and all reagents can be made available from the corresponding authors upon request. Source data have been deposited in NCBI’s Gene Expression Omnibus and are accessible through GEO series accession number GSE234396.

## MATERIALS AND METHODS

### Cell culture

HEK293T cell lines were maintained in dMEM medium (Gibco) supplemented with 10% fetal bovine serum and 1% penicillin/streptomycin. H1 human ESC lines (WA01, WiCell) were maintained in mTeSR plus (STEMCELL) on Matrigel (BD Biosciences) and passaged using TryplE in single cells. NPCs were generated from H1 hESC using STEMdiff Neural Induction Medium (STEMCELL) using the embryoid body (EB) protocol and maintained in STEMdiff Neural Progenitor Medium (STEMCELL) under non-differentiating conditions or cryopreserved using STEMdiff Neural Progenitor Freezing Medium (STEMCELL).

All cells tested negative for mycoplasma.

### Lentivirus production and stable gene expression

HEK293T cells overexpressing HA-tagged constructs were generated as described in de Tribolet-Hardy et al., 2023^1^. In short, cDNAs for ZNF274 constructs were codon-optimized, synthesized using the GeneArt service from ThermoFisher and cloned into the doxycycline inducible expression vector pTRE-3HA. Stable cell lines were generated using Lentivector transduction of HEK293T cells at an MOI of 5 or 10 as described on http://tronolab.epfl.ch and selected with puromycin (5 µg/ml) or hygromycin (200 µg/ml).

ZNF274 expression was induced with 1µg/ml doxycycline and verified via western blot with an anti-HA antibody (1:1000, ref.12013819001, Roche) and anti-beta actin antibody (1:1000, ref. ab20272, Abcam).

### CRISPR-Cas9 editing

*ZNF274* was deleted in HEK293T and H1 hESC cells using pX459 CRISPR-Cas9 system (addgene #62988). sgRNAs upstream and downstream of *ZNF274* were cloned into a Cas9-PuroR construct. Puromycin-resistant cells were selected after transfection with the two sgRNAs to increase the probability for a direct deletion.

The sequences of the guides are: for_2A (CACCCAAGCGGCGACACACGTTGC), rev_2A (AAACGCAACGTGTGTCGCCGCTTG), for_2B (CACCCAAACGTCCACCCTAGTTCT), rev_2B (AAACAGAACTAGGGTGGACGTTTG).

For HEK293T, 400’000 cells were transfected for 24h using Lipofectamine 300 (Mirus Bio, ref. MIR2300) and 2µg of sgRNAs, selected with 2µg/ml puromycin and single-cell sorted with flow cytometry in 96 well plate for screening.

For H1 hESC, 400’000 cells were transfected for 48h using TransIT-LT1 (Thermo Fisher, ref. L3000008) and 2.5 µg of sgRNAs, selected with 0.5 µg/ml puromycin and plated at low density in mTeSR Plus. Single colonies were picked and seeded in 96 well plate for screening.

Effective knockout of *ZNF274* was validated by PCR on genomic DNA and RT-qPCR on RNA transcribed into complementary DNA (cDNA) using the following primers:

- PCR on genomic DNA:
  - wild-type allele set 1 (ATCCAGGCCCTATATGCTGAA, GAGCGTGCCTTAGGCTGTA);
  - wild-type allele set 2 (GACAAGACCATACTGGGACC, CACGCAACTTTTGGGGAAGG);
  - deletion (GACAAGACCATACTGGGACC, AGGCGAGACGCTGATTGGAT).
- RT-qPCR on cDNA:
  - wild-type allele set 1 (ATCCAGGCCCTATATGCTGAA, GAGCGTGCCTTAGGCTGTA);
  - wild-type allele set 2 (TCAGCTTTCCAAACCAGATG, TGGAATGGTGTCTTGAGGAA).

### Chromatin immunoprecipitation

ChIP was performed as described previously^2^. Briefly, cells were cross-linked for 10 minutes at room temperature in PBS + 1% methanol-free formaldehyde solution, followed by quenching with 250 mM Tris pH 8. Nuclei were sonicated with Covaris E220 sonicatior (settings: 5 % duty, 200 cycle, 140 PIP, 20 min) to yield genomic DNA fragments with a bulk size 200-500bp. IP was performed overnight at 4° C using 5 μg antibody (anti-H3K9me3 Diagenode, ref. C15410056; anti-HA.11 BioLegend ref: 901503; anti-H3K4me3 Cell Signaling, ref. 9751S; anti-H3K36me3 Diagenode, ref. C15410192; anti-CTCF Millipore, ref. 07-729; anti-Rad21 Abcam, ref. ab993) coupled to 50µl Protein G Dynabeads (Invitrogen ref: 10009D). Final DNA was purified with QIAGEN Elute Column. Up to 10 ng of material for both total inputs and chromatin immunoprecipitated samples were used for library preparation. After end-repair and A-tailing, Illumina IDT indexes were ligated to the samples. Libraries were size-selected using Ampure XP beads (Beckman Coulter), quality-checked on a Bioanalyzer DNA high sensitivity chip (Agilent) and quantified with a Qubit dsDNA HS assay. Libraries were sequenced with 75 bp paired end on the NextSeq 500 (Illumina)

### Cut&Tag

CuT&Tag was performed as described previously (Kaya-Okur et al, 2019). For each mark, 200’000 cells were used per sample using the anti-H3K9me3 primary antibody (Active Motif, AB_2532132). A homemade purified ProtA-Tn5 protein (3XFlag-pA-Tn5-Fl, addgene #124601) was produced and coupled with MEDS oligos by the Protein Production and Purification of EPFL, as previously described (Bryson T, Henikoff S. 3XFlag-PATn5 Protein Purification and MEDS-Loading; 2019. doi:10.17504/protocols.io.8yrhxv6). Purified recombinant protein was used at a final concentration of 700 ng/uL. Libraries were sequenced with 75 bp paired end on the NextSeq 500 (Illumina).

### ProtA-TurboID

ProtA-TurboID experiments were performed as described previously^3^. A homemade purified ProtA-TurboID protein (Pk19 ProtA-Turbo plasmid) was produced by the Protein Production and Purification of EPFL. Briefly, HEK293T cells were cultured in 15 cm dishes and induced for 3 days with 1µg/ml doxycycline. Cells were fixed in 4 ml of PBS +4 % PFA for 15 min at RT and washed twice with PBS. Cell pellets were incubated for 10 min on ice in hypotonic lysis buffer (10 mM Tris pH 7.5, 10 mM NaCl, 3 mM MgCl2, 0.3 % NP40, and 10 % glycerol), then centrifuged and washed to isolate nuclei. Nuclei were permeabilized with 0.3% Triton-X100 for 10 min. Non-specific-binding sites were blocked with blocking solution (PBS + 0.3 % Triton-X100 + 3 % BSA) for 30 min in a rotation wheel at RT. Samples were then incubated with 3 μg of anti-HA antibody (Abcam, ref. ab9110) diluted in 300 μl of blocking solution for one hour at RT. After washing, nuclei were incubated with 3.5 μg of ProtA-Turbo enzyme in 300 μl blocking solution for one hour at RT. Next, samples were incubated with 300 μl of biotin reaction buffer (5 mM MgCl2, 5 μM Biotin, 1mM ATP in PBS) in a thermo shaker at 1000 rpm for 10 min at 37 °C and finally lysed overnight at 4° C in 300 μl of RIPA buffer (50 Mm Tris pH 7.8, 150 mM NaCl, 0.5% sodium deoxycholate, 0.1% SDS, 1% NP40). The following day, samples were sonicated using a Branson LPe Sonicator (3 cycles of 20 s at 30 % amplitude) and decrosslinked for 1 h at 95 °C after addition of SDS to a final concentration of 1 %. After an additional sonication cycle, samples were centrifuged and supernatant was incubated with 12.5 μl streptavidin sepharose high-performance beads (15511301, Cytiva) on a rotation wheel for 2 h at 4° C. Prior to use, streptavidin beads were chemically modified to be protease-resistant as described in Rafiee et al., 2020^4^. After streptavidin pulldown, beads were washed 5 times with RIPA buffer, 5 times with PBS and eluted in elution buffer (2 M Urea, 10 mM DTT, 100 mM Tris pH 8). After incubation at 1500 rpm for 20 min at RT, 50 mM iodoacetamide was added and samples were incubated at 1500 rpm in the dark for 10 min at RT. 2.5 μl of trypsin was then added to each sample, followed by incubation at 1500 rpm for 2 h at RT. The eluates were collected and 1μl of fresh trypsin was added for overnight incubation at RT. The following day, peptides were acidified, purified on C18 Stagetips subjected to LC-MS.

### Coimmunoprecipitations

Coimmunoprecipitations were performed as described previously^5^. HEK293T cells cultured in 10 cm dishes were transfected for 24 h in 1 µg/ml doxycycline with plasmids carrying HA-tagged constructs. Cells were washed three times PBS, lysed 30 min at 4°C in 500 µl IP buffer (0.5 % NP-40, 500 mM Tris pH 7.4, 20 mM EDTA, 10 mM NaF, 2 mM benzamidine) and centrifuged 3 min at 5000 rpm. A fifth of the lysate was taken as WCE control and the rest was added to 25 µl of appropriate anti-Flag sepharose beads or anti-HA magnetic beads for overnight incubation at 4° C. After washing, proteins were eluted off the beads in 50 µl of 2X loading buffer and boiled for 10 min. 15 µl of IPed proteins and 10 µl of WCE were submitted to SDS-PAGE and analyzed by immunoblotting using anti-HA antibody (1:250, ref.12013819001, Roche), anti-Flag antibody (1:500, ref.A8592, Sigma-Aldrich) and anti-Kap1 (1:250, ref. ab10843, Abcam) antibodies.

Experiments were repeated three times and representative blots are shown in the figures.

### GFP reporter assay

*ZNF274* KO HEK293T cells were transduced with pRLL-PGC-GFP or pRLL-SNP-GFP constructs at MOI 0.5 and GFP+ cells were isolated by flow cytometry. GFP+ cells were subjected to a second transduction with MOI 10 of pTRE-ZNF274 inducible constructs and selected for puromycin resistance. GFP+ HEK293T transduced with pTRE-ZNF274 constructs and control cells were treated or not with doxycycline and GFP signal was read by flow cytometry.

### Immunofluorescence

HEK293T cell lines were grown on coverslips covered with 0.01 % poly-L-lysine (Sigma-Aldrich) and fixed for 15-20 min in a 4% methanol-free formaldehyde solution. Cells were washed three times with PBS, permeabilized with 0.5 % Triton X-100 for 20 min and blocked with 1 % BSA for 30 min. Cells were incubated with anti-HA antibody (1:1000, ref. ab9110, Abcam) and anti-B23 (1:500, ref. B0556, Sigma-Aldrich) in 1 % BSA overnight at 4° C or 1 h at RT. Samples were washed three times with PBS and incubated with Alexa 647-conjugated anti-mouse antibody (1:1000), A488 anti-rabbit antibody (1:1000) and DAPI (1:10’000) in 1 % BSA for 1 h at RT. Three final washes were performed before mounting the slides in Vectashield (Vector Laboratories). Images were acquired on a Leica-SP8 confocal microscope using the oil objective HC PL APO 63x/1.40 and analyzed with ImageJ software (version 2.9.0/1.53t).

### DNA-FISH

Two clones for wild-type and *ZNF274* KO background were karyotyped prior to use for DNA-FISH using the CytoScan Array to ensure consistent ploidy.

HEK293T cells were grown on coverslips covered with 0.01 % poly-L-lysine (Sigma-Aldrich). Cells were fixed for 10 min in a 3 % methanol-free formaldehyde solution and permeabilized for 5 min in ice-cold CSK buffer (10 mM PIPES; 300 mM sucrose; 100 mM NaCl; 3 mM MgCl_2_; pH 6.8) supplemented with 0.5 % Triton X-100 and 1 mM EGTA. After RNase treatment for 1 h at 37°C with 100 µg/ml RNase A and stored at −20° C or used directly for DNA-FISH.

DNA-FISH probes were obtained after nick translation of BAC constructs (RP11-45IJN, RP5-915N17) purified using the Large Construct kit (Qiagen): 1 μg of purified DNA was labelled for 3 h at 16° C with fluorescent dUTPs (SEEBRIGHT Orange 552 dUTP and SEEBRIGHT Green 496 dUTP, Enzo Life Sciences). 100 ng of probes were supplemented with 5 μg of Cot-I DNA (Invitrogen) and 10 μg of Sheared Salmon Sperm DNA (Invitrogen). After precipitation, the probes were resuspended in deionized formamide, denatured for 7 min at 75° C and further incubated for 15 min at 37° C. Probes were mixed with an equal volume of 2X Hybridization Buffer (4X SSC, 20 % dextran sulfate, 2 mg/ml BSA). Next, coverslips were denatured for 10 min at 80° C in 70 % formamide/2X SSC, dehydrated in 80-100 % ethanol washes and incubated with the hybridization mix at 37° C overnight in a humid chamber.

The following day, coverslips were washed for 4 min at 42° C three times with 50 % formaldehyde/2X SSC (pH 7.2) and three times with 2X SSC. The, IF was performed at this step using anti-Nucleolin (1:1000, ref. Ab22758, Abcam) and coverslips were mounted in Vectashield (Vector Laboratories). Images were acquired on a Leica DM5500 upright motorized microscope, using the oil objective HCX PL APO 63X/1.40 and with a z-stacks collected at 0.2 µm steps for each channel.

Images were processed by ImageJ (version 2.9.0/1.53t) and deconvolved using Huygens Remote Manager (version 3.7). We used the ImageJ plugin for StarDist to detect nuclei from individual cells based on DAPI staining^6^. The Distance Analysis (DiAna) plugin was used to count the number of DNA-FISH foci per cell and calculate the distance between the DNA-FISH signal and the nucleolar marker immunofluorescence signal^7^. Cells with three DNA-FISH spots for a given DNA loci were manually identified and the number of cells with at least one contact with the nucleolus was visually confirmed.

DNA-FISH experiments have been performed twice and the percentages displayed are computed from pooled values of biological replicates.

### Hi-C

The Hi-C libraries were prepared using Arima-HiC (Arima Genomics), following the manufacturer’s protocol. For all samples, either 1 million or 5 million cells were fixed in 1 % methanol-free formaldehyde solution for 10 min at RT and the reaction was quenched for 5 min with glycine to a final concentration of 125 mM. Libraries were sequenced with 100 bp paired end on the NovaSeq 6000 (Illumina).

### UMI-4C

UMI-4C experiments were performed as described previously^8^. Briefly, 5 million cells were fixed in 1 % methanol-free formaldehyde solution for 10 min at RT and the reaction was quenched for 5 min with glycine to a final concentration of 125 mM.

Cells were lysed in ice-cold Lysis Buffer (10 mM Tris HCl pH 8, 10 mM NaCl, 0.2 % NP-40), chromatin was digested with 800U DpnII (NEB), and digested ends were marked with biotin (ref. 19524016, Life Technologies) prior to ligation.

Universal Primer 2 was used in all the reaction as the reverse primer (CAAGCAGAAGACGGCATACGA). The following upstream and downstream primers were used for PCR amplification:

- DS1_Peak2A_PCDH: AATGATACGGCGACCACCGAGATCTACACTCTTTCCCTACACGACGCTCTTCCGATCTTGTGTGATGCCATGTGTGTT
- US1_Peak2A_PCDH: CAAGGAACACATAAAGGATTCC
- DS3_Peak2A_PCDH: AATGATACGGCGACCACCGAGATCTACACTCTTTCCCTACACGACGCTCTTCCGATCTGTGATGCCATGTGTGTTTTTAAC
- US3_Peak2A_PCDH: AGCCCACTGTGAGTTCTTACAT
- DS1_Peak1_PCDH: AATGATACGGCGACCACCGAGATCTACACTCTTTCCCTACACGACGCTCTTCCGATCTTATCCACTACTGGTCATCTCA
- US1_Peak1_PCDH: CATCTATTGTGTTTTTCACAGTATG
- DS2_Peak1_PCDH: AATGATACGGCGACCACCGAGATCTACACTCTTTCCCTACACGACGCTCTTCCGATCTACTTATACATTTGTTCTGTT
- US2_Peak1_PCDH: AGAGCCAGAAACTATTATTGTA
- DS1_HS5-1: AATGATACGGCGACCACCGAGATCTACACTCTTTCCCTACACGACGCTCTTCCGATCTCATTCCCCGTTGTTTTGGCG
- US1_HS5-1: GTGGGTTAAATTAAGCCGAGTT
- DS2_HS5-1: AATGATACGGCGACCACCGAGATCTACACTCTTTCCCTACACGACGCTCTTCCGATCTCTGAAAAATGCAGTCGACTCGC
- US2_HS5-1: TTAATTTCCCAGGGGCAGAG
- DS1_HS16-20: AATGATACGGCGACCACCGAGATCTACACTCTTTCCCTACACGACGCTCTTCCGATCTTTGTGAGGTCAGAGACTAGGGA
- US1_HS16-20: TGCATTTTTCTTTTCGGCCAC
- DS2_HS16-20: AATGATACGGCGACCACCGAGATCTACACTCTTTCCCTACACGACGCTCTTCCGATCTACCGCCATCTCTGTTACTGA
- US2_HS16-20: GTGGAAGCCAGGAAAGGGTG

Libraries were sequenced with 75 bp paired end on the NextSeq 500 (Illumina).

### EM-seq

200 nanograms of DNA from each cell line were processed following the NEB EM-Seq protocol (E7120S). After library preparation, the DNA was quantified and sequenced with 100 bp paired end on an Illumina NovaSeq to obtain ∼100 million reads/sample.

### 10x Genomics 5’ single-cell RNA-seq

Wild-type and *ZNF274* KO NPCs were detached using Accutase (07922, STEMCELL), washed in PBS and immmediately processed for 10x library preparation using the Single Cell 5’ v2 Gene Expression Library. All libraries were sequenced on an Illumina NovaSeq. For each condition, we aimed to collect ∼4,000 cells per sample.

## Data analyses

### ChIP-seq and Cut&Tag

Raw reads were mapped to the human genome (hg19) using the short read aligner program Bowtie2, with the sensitive local mode. Before peak calling, bams were filtered to remove low quality reads (mapq < 10) and then run through the Bioconductor package GreyListChIP (R package version 1.32.0) to mask artefacts regions. For ChIP-seq, peaks were called with MACS2 using the –bampe option for the transcription factors and EPIC was used for the histone marks^9,10^. High quality peaks were kept via thresholding for score > 50 for MACS2 and FDR<0.01 for EPIC and merged using the R package DiffBind so that only peaks that occur in at least 2 peak sets will be kept and merged into consensus peaks. For Cut&Tags, peak calling was done using SEACR with stringent parameters^11^. ChIP-seq derived motifs were found using the online RSAT tool with default parameters. BEDtools was used to perform overlap and genomic analysis alongside the R package Chipseeker for annotation and Deeptools for coverage plot^12–15^. Differential binding analysis was done in R. After read quantification on peaks using BEDtools multicov -q 10 on merged peaks, a countable was produced and normalized for sequencing depth using the TMM method as implemented in the limma package of Bioconductor^16^. A moderated t-test from the limma package was used to test significance. P-values were corrected for multiple testing using the Benjamini-Hochberg’s method^17^. Regions with absolute foldchange bigger than 2 and p-adjusted smaller than 0.05 were considered significant.

### RNA-seq

Reads were mapped to the human (hg19) genome using hisat2 with parameters -k 5 --seed 42 -p 7^18^. For stranded data, the --rna-strandness RF parameters was added. Counts on genes and TEs were generated using featureCounts^19^. To avoid read assignation ambiguity between genes and TEs, a gtf file containing both was provided to featureCounts. For repetitive sequences, an in-house curated version of the Repbase database was used (fragmented LTR and internal segments belonging to a single integrant were merged, as in Turelli et al^20^. Only uniquely mapped reads were used for counting on genes and TEs. Finally, the features for which the add up of reads across all samples were lower than the number of samples were discarded from the analysis. Normalization for sequencing depth was done for both genes and TEs using the TMM method as implemented in the limma package of Bioconductor and using the counts on genes as library size. Differential gene expression analysis was performed using voom as it has been implemented in the limma package of Bioconductor. A gene was considered to be differentially expressed when the fold change between groups was bigger than 2 and the p-adjusted was smaller than 0.05. A moderated t-test (as implemented in the limma package of R) was used to test significance. P-values were corrected for multiple testing using the Benjamini-Hochberg’s method^17^.

### UMI4C

UMI4C data were preprocessed and analyzed with the R package umi4cPackage (with default parameters) using on a custom genome including the rDNA sequence as additional chromosome (NCBI U13369.1)^8^. For comparing the KO to the WT, the KO sample was normalized to WT and the domainogram shows the fold change in mean contact intensity between the two conditions. Fold changes at specific regions and p-values were computed via a Chi-square test and error bars were estimated from the binomial s.d.

### Hi-C

Raw reads were processed through HiC-Pro v 3.1.0 pipeline^21^. Mapping parameters were kept to default to map on a custom genome including the rDNA sequence as additional chromosome (NCBI U13369.1). The digestion parameters were set according to the restriction enzyme (GATCGATC, GANTGATC, GANTANTC, GATCANTC) and the minimum and maximum insert size set to 100 and 1400 respectively. The contact maps were generated for 20kb and 100kb then normalized using the ICE methods. Analysis was done using the R package GENOVA^22^.

### Turbo-ID

LFQs were generated using the MaxQuant pipeline and then normalized using the variance stabilization normalization (VSN) as implemented in the DEP R package^23,24^. Missing values were imputed differently depending if the data was missing at random (MAR) or not (MNAR). For MAR, the k-nearest neighbors approach was used with option rowmax=0.9. For MNAR values, the imputation was performed drawing from a gaussian distribution using the minProb option with parameter q=0.1. For the contrasts between conditions, the threshold of FDR=0.05 and log2 foldchange 1.5 were chosen. Gene set enrichments were computed using the hyperGTest from the GOstats R package with pvalue cutoff of 0.05^25^.

### EM-seq

EM-seq raw data were processed by the Msuite2 v.2.2.0 pipeline with parameters −3 -m BS and using hisat2 as mapper on the hg19 version of the human genome with rDNA added as extra chromosome^26^. The resulting mean methylation calls across replicates were aggregated by regions of interest and the 75^th^ percentile of all CpG methylation in the region were computed as summary statistic for final analysis. Plotting was performed in python3 using seaborn package and the statistical significance assessed by Wilcoxon test using the scipy python package^27,28^.

### 10X scRNAseq

10x data were processed by standard cellranger (v7.0.0) pipeline on the hg19 genome^29^. The count data was then loaded in Seurat v 4.3.0 for QC and analysis^30^. Cells with more than 200 features and less than 25% mitochondrial gene content were kept for further analysis. Seurat’s SCTransform was used to normalize the data and correct for mitochondrial percentage and total number of reads biases. UMAP were computed using 30 principal components on the transformed data in Seurat. For estimating the number of kZFP and PCDH genes detected per cell, a gene was considered detected as soon as harboring at least one read. Then, the Seurat FindClusters function was used with resolution 0.2 to infer cell clusters. 2 clusters were identified as outliers and correlated with high mitochondrial gene expression and were discarded from the rest of the analysis. For graphical representation, scCustomize R package was used^31^.

## Notes

### Competing Interest Statement

M.B. and D.T. are inventors on an international patent application submitted by the Ecole Polytechnique Federale de Lausanne for the exploitation of the SCAN domain in genome-editing technologies (European Patent Application EP 23183405.2).
The remaining authors declare no competing interests.

